# Frame Effects Across Space and Time

**DOI:** 10.1101/2025.10.24.684468

**Authors:** Bernard Marius ’t Hart, Patrick Cavanagh

**Author notes:** Corresponding author: Marius ‘t Hart, Centre for Vision Research, York University, Toronto, ON M3J1P3, Canada.

## Abstract

When two probes are flashed at different times within a moving frame they can be perceived as dramatically separated from each other even though they are at the same location in the display. This effect suggests that we perceive object position relative to the surrounding frame even when it is moving (Özkan et al., 2021). Here, 8 experiments reveal new properties of this frame effect. First, the influence of the frame on the perceived probe positions extends beyond its bounding contours by several degrees of visual angle, both in the direction of the frame’s motion and orthogonal to it. It is also undiminished when the probes and the frame are in different depth planes. However, the influence of the frame’s motion shows no extension in time – there is no effect on probes presented after the frame is removed and none retroactively before the frame appears either. The frame effect is also driven primarily by the displacement of the frame, not by its motion signals: the effect is stronger for moving bounded frames compared to moving, unbounded random-dot textures. When the bounded region has an internal texture that moves with or against the frame’s motion or remains static, it is the displacement of the frame that produces the perceived position shifts of the probes, while the effect of the internal motion is mostly suppressed. The frame’s influence is unaffected by whether the motion is self-initiated or not and does not reduce in strength across 2 hours of testing.

## Introduction

Correctly perceiving the location of objects is important when interacting with them but this can be complicated when the object is part of a moving group or surface. In a typical example (Özkan et al., 2021; Cavanagh et al., 2022; Shams et al., 2024), a square frame moves continuously left and right and two probes are flashed at the same horizontal location at the center of the frame’s path, once when the surrounding frame is at the right extreme of its motion path and then again when it is at the left extreme. In this case, even though the probes are physically aligned on the screen, they can appear offset from each other by as much as the distance the frame moves. That is, the locations of the probes are perceived relative to the moving frame, one on the left of the frame, the other on the right, rather than in absolute coordinates (Fig. 1). A previous study (Cavanagh et al., 2022) suggested that the position shift relies on the probes belonging to or grouping with the frame. Critically, these frame-relative locations are perceived as if in a “virtual” frame, stabilized at the midpoint of its path. The virtual frame that serves as the reference for the probe locations is itself not seen, whereas the physical frame does appear to move, although through a shorter distance than the actual path (Özkan et al., 2021).

**Figure 1.**
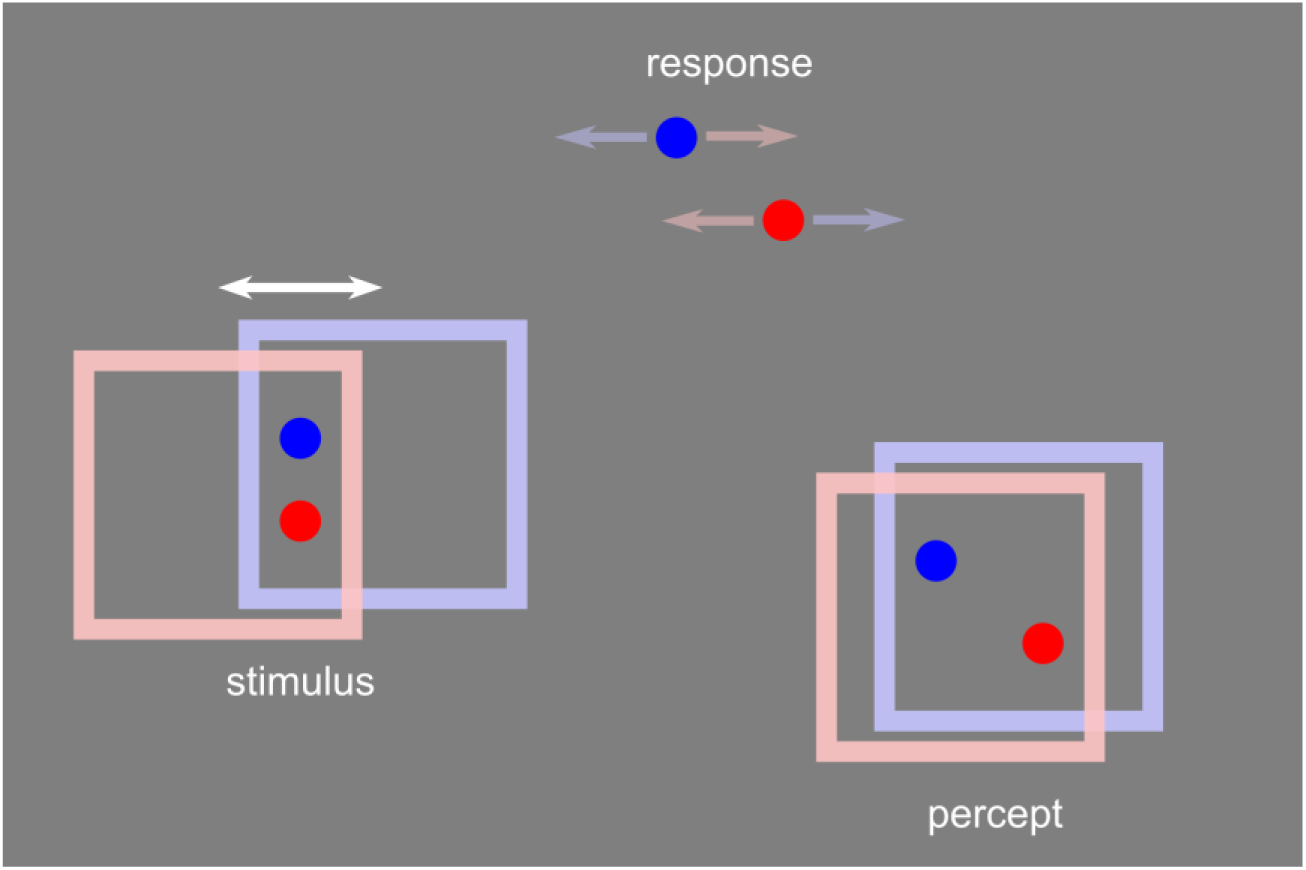
The Frame Effect. Two probes are briefly flashed in a frame moving left and right. The blue probe flashes when the frame is at its rightmost end of travel and the red when the frame is at the left end of its travel. Although aligned vertically on the screen, the flashed probes are perceived at their locations relative to the frame at the time they flashed (blue on the left, red on the right), as if the frame were almost stationary. Participants report their percept by adjusting two reference markers such that the angle of the markers matches the perceived angle between the flashed probes. The frames are colored here to indicate the frame location during the presentation of the probe of the corresponding color. In the experiments, the frames were always white. The frames are also vertically offset here for clarity whereas in the experiments, the motion was purely horizontal.

**Movie 1.**
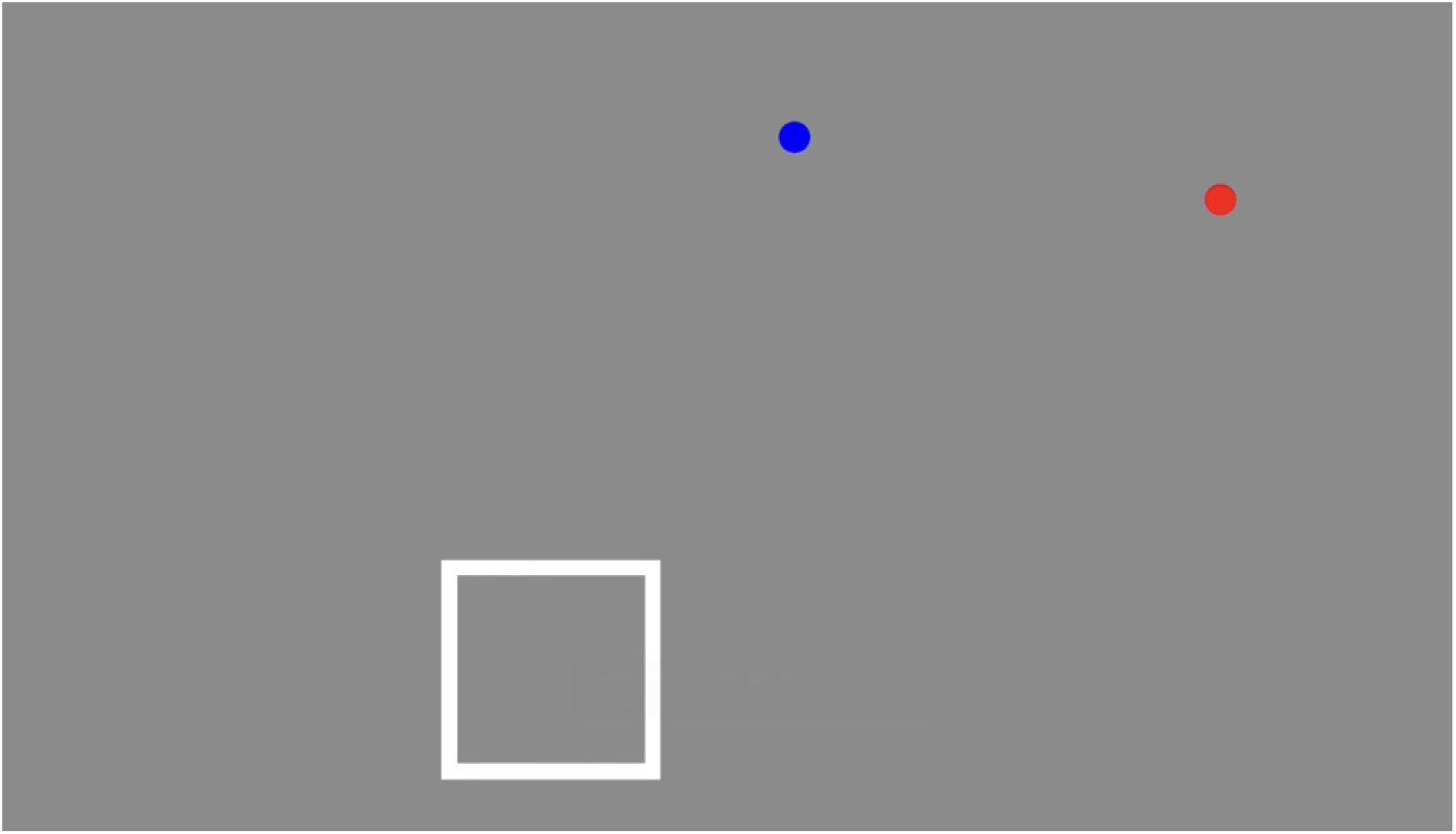
Measuring the frame effect. Participants adjusted the two top dots so their angle matched the perceived angle between the blue and red dots that flashed inside the moving frame. The sequence repeated until the participant responded. Use this link to open the movie in a browser window: https://cavlab.net/Demos/FEAST/#0/.

Frame effects have a long history. They have been shown to influence the perception of an object’s motion (Johansson, 1950) or our own (Warren, 1895), to change what direction we judge to be “up” (Asch & Witkin, 1948; Morgan, Grant, Melmoth & Solomon, 2015) or straight ahead (Roelofs, 1935; Matin & Fox, 1989). Here we consider the effect of a moving frame on the perception of position where Duncker (1929) provided the first compelling demonstrations. He reported that a static probe presented within a moving frame appeared to move in the direction opposite of the frame. In this case, the induced motion is clear but there is only a small offset in the perceived position of the two ends of the induced travel (MacLeod et al, 2024). Wallach et al., (1978) tested the effect of a moving frame on a probe that was not static but instead moved orthogonal to the frame’s movement. This increased the perceived spatial offset between the two ends of its path (e.g., Anstis & Cavanagh, 2024). However, if the probes are neither static nor in continuous motion but flashed at each end of the frame’s travel, the apparent spatial offset becomes dramatically larger (as much as 6 times larger, Anstis & Cavanagh, 2024). Duncker (1929; chapter 4) first reported this effect with flashed probes using a frame in apparent motion, as did Wong and Mack (1981). Recent studies have demonstrated that this large separation between the flashed probes is seen with either continuous and apparent motion of the frame (Özkan et al., 2021; Cavanagh et al., 2022; Chung & Stormer, 2023). These studies showed that the frame effect is quite tolerant of variations in the shape of the frame and its motion path, but they did not address the spatial or temporal extents over which the frame influenced flashed probes. The eight experiments here vary the spatial and temporal offsets between the probes and the frame; they also examine the effects of separate motions of the frame and the texture within the frame, the effect of self-generated frame motions and, finally, the effects of practice.

## General Methods

### Participants

14 unpaid volunteers were recruited as participants, including the first author (age: 20-45, avg 27.9, females: 4, left-handed: 1). All had normal or corrected-to-normal vision and provided prior, written informed consent. All procedures were approved by the York Human Participants Research Committee.

### Apparatus

Stimuli were presented on a Dell E2009Wt monitor (1680x1050 pixels; 60Hz; 43.3x27.1 cm) which was kept at 60 cm distance from participants eyes using a chin rest. The screen’s gamma function was linearized and the background set at 50% luminance gray. Experiments were controlled by software in PsychoPy (Peirce et al., 2019) on a Dell Latitude E6530 with an NVidia NVS 5200M graphics card. In Experiment 6 (self-moved frames), the participants moved the frame by themselves using a stylus on a drawing tablet (Intuos Pro Large). In Experiment 2, red/cyan filter glasses were worn by the participants to create an impression of depth between the probes and the frame.

### General Stimuli

The standard frame was a 7x7 dva square, consisting of a white (100% of the monitor’s luminance) edge of 0.5 dva width (leaving a 6x6 dva opening). The standard frame motion spanned 4 dva. The two probes were blue (top) and red (bottom) circles with a diameter of 1 dva positioned 1 dva above and below the middle of the frame path. The frame stopped moving for 133.3 ms at the extremes of its horizontal path during which time the probes were presented, ensuring that the frame was never in motion when the probes appeared. The probes were presented for 66.7 ms centered in the middle of the motion pause. Variations from these values are described in each experiment when present.

For all trials, participants were given two continuously visible reference dots, above and to the right of the frame and probes, whose position could be controlled with a mouse. To avoid the influence of peripheral effects on perceived size (Newsome, 1972), participants were instructed to position the reference dots such that their slant matched that of their percept, rather than matching the extent of the separation. The stimulus cycled through its motion until the participant responded. Participants were free to look anywhere, at the stimulus or the reference dots, during most trials (exception in exp. 5). The perceived offset between the two flashed probes was recorded as the horizontal distance between the two reference dots. The frame motion was randomly flipped left-right on each trial so that the effect’s direction was not predictable.

Experiments were divided into blocks, each of which repeated the same trials, but in a shuffled order. The number of blocks per experiment is described in each method section. The order of experiments was shuffled across participants as well. For each participant, the seed of the random number generator used to generate shuffled conditions and tasks was set using the participant’s ID.

Data were processed and statistically analyzed in R 4.5.1 (R Core Team, 2025). All data and scripts are available on https://osf.io/ufzsq and https://github.com/thartbm/FrameEffects.

## Effects in space

The frame effect requires that the probes group together with the moving frame as opposed to the static background (Cavanagh et al., 2022). Does this grouping occur only when the probes are inside the frame, or does it extend some distance outside it as well? The two tasks here tested the spatial relationship between the flashed probes and the frame. The first tested the effect of horizontal and vertical offsets, as well as the size of the frame while the second tested the effect of offsets in depth between the probes and the frame.

### Exp 1. Frame spatial offset and size

The offset between the probe locations and the frame was varied to see if effect of the frame’s motion extends outside the frame and whether it does so similarly in the direction of motion and the direction orthogonal to the motion (Figure 2 and Movie 2). The frame’s effect may also depend on the distance of the probe to the frame contour or on the distance to the frame center. This was taken into account by varying the frame’s size.

**Figure 2.**
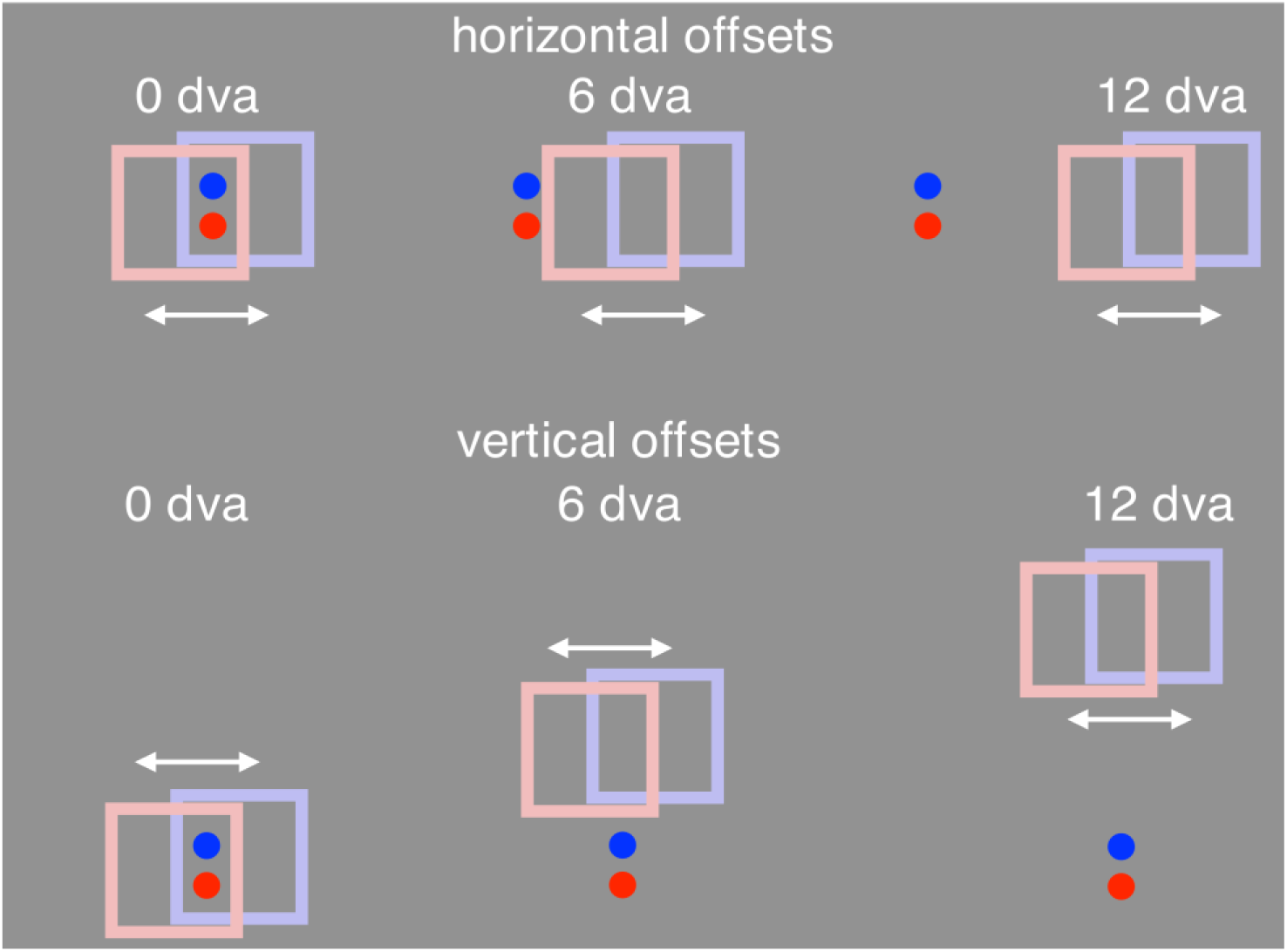
Spatial Offsets. The frame’s path was shifted either horizontally (top row) or vertically (bottom row) by one of 5 values (0, 3, 6, 9, or 12 dva). The locations of the probes and response markers remained unchanged on the screen for the different frame offsets. The offsets were equally often left or right (horizontal) or up or down (vertical). The offsets depicted here are for frames of 7x7 dva size whereas in the experiment the frame could be one of three sizes: 4x4, 7x7 or 10x10 dva. The frame colors and slight vertical shifts here are for clarity only.

#### Methods

In the first task, the frame was offset from the probes by 0, 3, 6, 9 or 12 dva, either horizontally (to the right along the direction of motion of the frame) or vertically (upward, orthogonal to the direction of motion of the frame). For each frame offset, the frame had one of three sizes, 4x4, 7x7, or 10x10 dva (in each case with a 0.5 dva wide outline going inward). At the smallest size, the probes overlapped the edge of the frame, and this also happened at some of the offsets for the larger frames. The duration of the 4 dva motion was 250 ms in each direction with a 100 ms pause at each end. There were 30 main conditions (3 frame-sizes x 5 frame offsets and 2 offset directions) presented in 5 blocks. Interspersed within these offset and size trials were 4 control trials in each block for testing the effect of time-on-task on the strength of the perceived offset. For these trials there was no vertical or horizontal offset and the 7x7 dva frame had 0.8, 1.6, 2.4 or 3.2 dva of motion (keeping frame motion duration constant at 250 ms). Similar control trials were also embedded in Experiments 3, 4, 5 and 6. There were 34 trials per block and 170 trials total.

**Movie 2.**
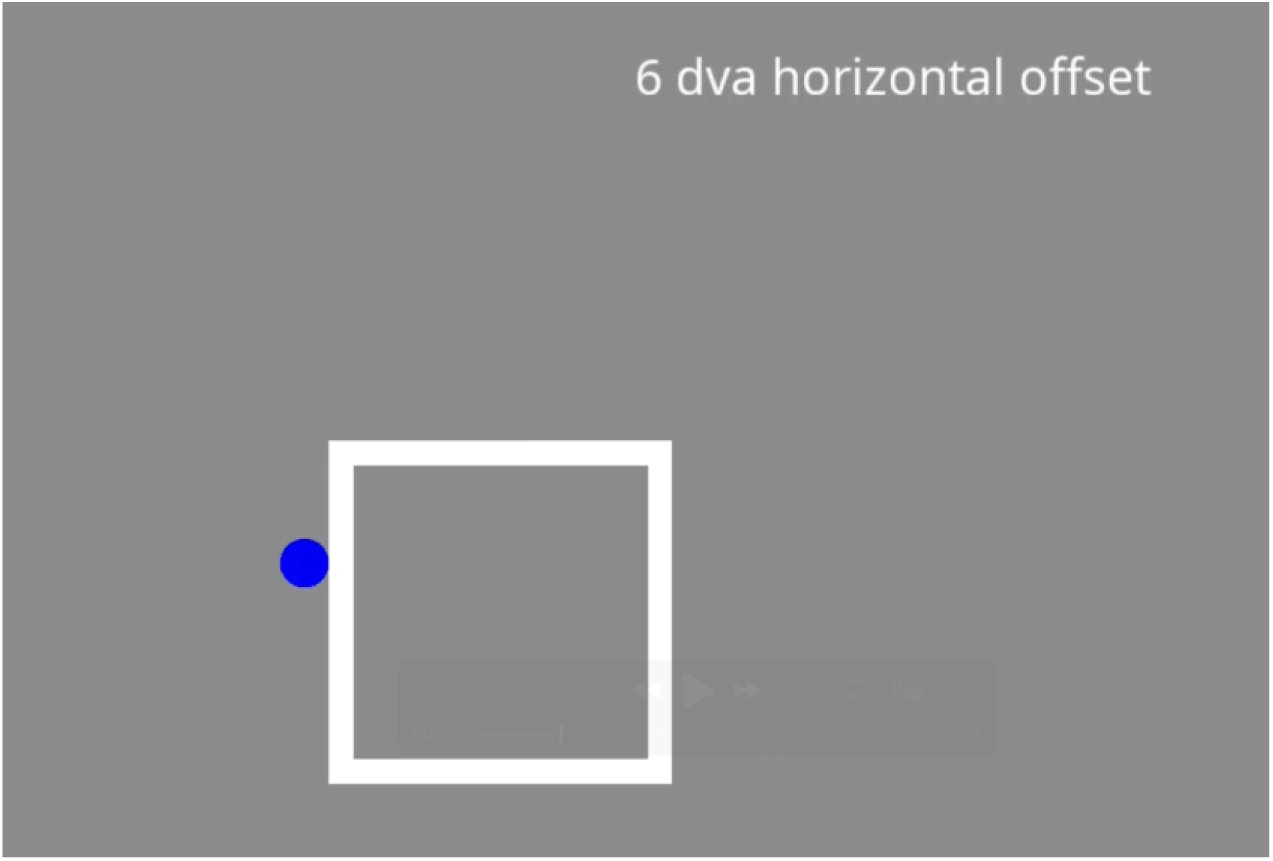
Varying spatial offsets and frame size. The adjustable dots in the top right corner are not shown in this demonstration movie but were present in the experiment. Use this link to open the movie in a browser window: https://cavlab.net/Demos/FEAST/#2/.

#### Results and Discussion

For both the horizontal and vertical offset trials, the analysis was based on the medians of the reported probe offsets across the 5 trials within each condition for each participant. Figure 3 shows the average of these medians across participants, along with a 95% confidence interval of the mean, assuming a *t*-distribution. The frame effect (the illusory separation between the two flashed probes) decreased following a sigmoid function as the distance between the frame and probes increased. This suggests that frame effect and the “grouping” of the frame and probes was strongest when the probes were within the frame, with the effect size ranging from 75% to even 100% of the 4 dva frame movement in these cases. The effect still held for probes outside the frame but dropped to little or no effect once the probes were 9 to 10 dva from the center of the frame’s path in either the horizontal or vertical direction.

**Figure 3.**
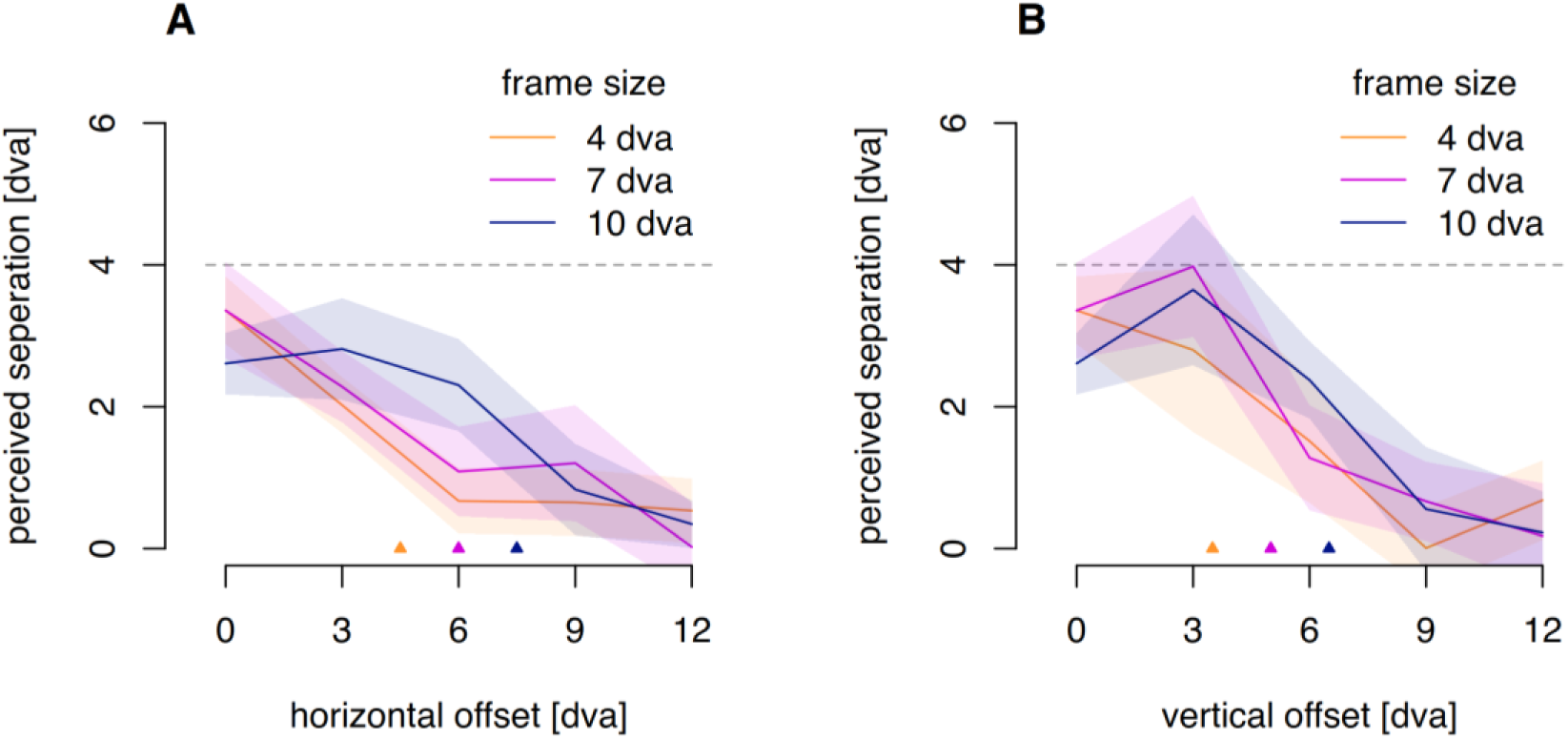
Frame size and frame offsets. The two probes were vertically aligned in the display. A. Perceived separation between the two probes for horizontal frame offsets and frame sizes. B. Perceived separation for vertical frame offsets and frame sizes. Colored lines indicate average reported probe separation for a given frame size as a function of frame offset, with the shaded region indicating a 95% confidence interval. The dashed horizontal line at 4 dva indicates the frame displacement amplitude. The small triangles along the horizontal axes in both panels indicate the offset beyond which the probes are always outside the frame.

A 3-way repeated measures ANOVA was run on the reported offsets, using frame size, frame offset distance and frame offset direction as factors. There was a main effect of offset distance (F(2.11, 27.44)=93.91, p<.001, η_G_^2^=.427), as well as a main effect of frame size (F(1.63,21.14)=7.98, p=.004, η_G_^2^=.033). These two factors also interacted (F(3.45, 44.86)=6.17, p<.001, η_G_^2^=.065). However, there was no effect of horizontal versus vertical offset direction (F(1,13)=3.95, p=0.68, η_G_^2^=.012) nor did offset direction interact with frame size (F(1.32,17.18)=3.22, p=0.81, η_G_^2^=.057). There were no other effects.

The offset of the frame mattered for the illusory separation seen between the two probes but the effect was similar for the vertical offset orthogonal to the frame’s motion and the horizontal offset parallel to the motion. These different directions of offset would affect the mean distance between the frame’s contours and the probes at the time of the flash differently, suggesting that the distance to the contour is not the primary factor determining the influence of the frame’s motion on the perceived probe position. It may instead be the distance of the probes from the mean location of the frame that determines the illusory shift. There was, nevertheless, an effect of the size of the frame which suggests there is an overall factor of size which modulates the frame effect – very small frames may not act as frames at all.

### Exp. 2 Depth offsets

In the second experiment, the probes and frame were placed at different depths to determine whether grouping of the probes and frame and the resulting frame effect depends on depth. The apparent depths produced by the disparity differences between the probes and the frame were about ±5 cm at a viewing distance of 60 cm. A separate task was run to verify that the depths for these disparities were correctly seen.

#### Methods

There were 4 conditions in this task. In the “same-plane” condition, both probes and the frame had zero disparity relative to the fixation (all stimuli in the same plane). In the “in-front” or “behind” conditions, the flashed probes had -0.5 dva disparity and the frame +0.5 dva (probes closer than the frame) or vice versa (frame closer than the probes). Finally, in “straddle” condition, the frame’s disparity was 0 dva, while the top probe had a +0.5 dva disparity and the bottom probe -0.5 dva disparity (the top probe in front of and the bottom behind the frame). There were 3 blocks with 20 trials in each (5 repetitions of each of the 4 conditions) for a total of 60 trials.

Before starting the task, participants adjusted the red and cyan colors of two separate stimuli, such that the visibility of the red stimulus through the cyan filter and the visibility of the cyan stimulus through the red filter were minimized as much as possible. They also performed a short test task, where they were shown two probe dots at the center of the screen, alternating one after the other without the frame (with the same size and relative vertical offset as the two flashed probes presented with the frame). Participants reported which stimulus (top or bottom) was in front. The top and bottom probes could have -0.5, 0 or 0.5 dva disparity for 6 possible combinations where the probes always differed by either 0.5 or 1.0 dva disparity, repeated 8 times for 48 trials total. One participant could not do this control depth task and did not run in the main task. Of the remaining 13 participants, 11 responded over 75% correct in this depth test (mean of 97% correct) and these participants were used for the analysis.

#### Results and Discussion

Figure 4 shows the reported separation between the two probes in the four arrangements of the frame and flashed probes in depth. The mean offset reported when the probes and frame were all in the same depth plane was 3.0 dva or 75% of the frame’s travel. This result is similar to the comparable setting found in Experiment 1 (for the 7x7 dva frame with no horizontal or vertical offset). The settings were also similar for the three other conditions that did have depth differences between the frame and probes. None of the means fell outside the 95% confidence interval of the others. This lack of significant difference across conditions was confirmed by a one-way, repeated-measures ANOVA on median percepts across the four stimulus types (F(2.04, 20.40)=0.98, p=.395, η_G_^2^=.017).

**Figure 4.**
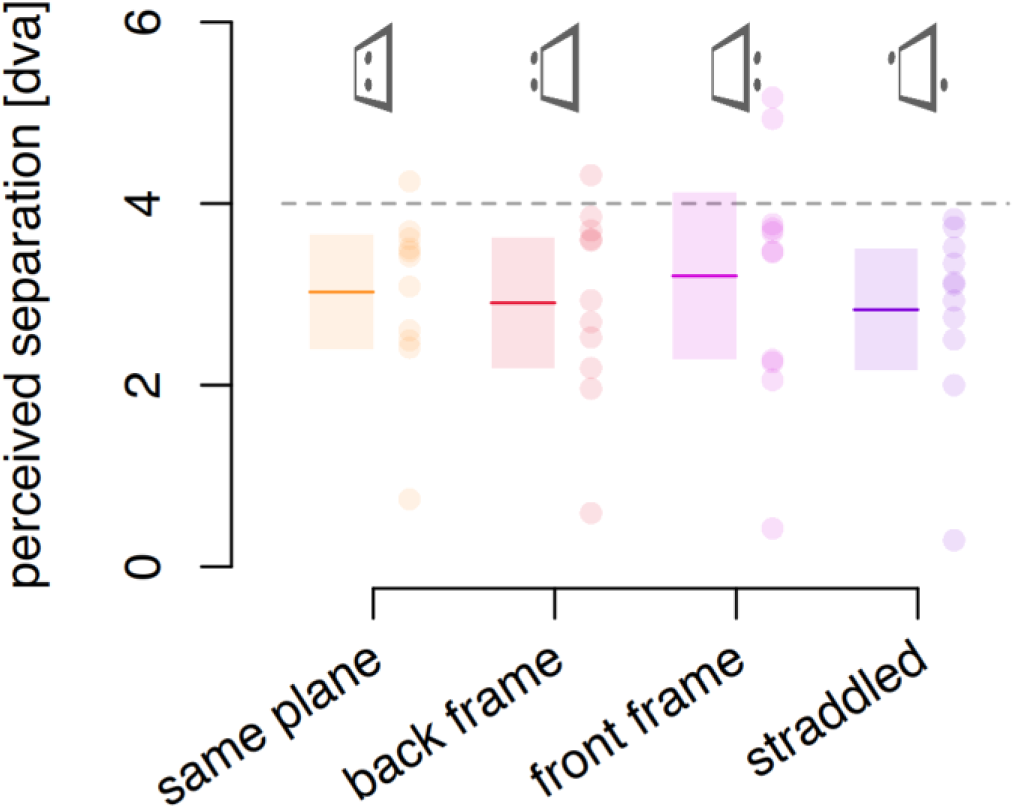
Effects of depth offsets between frame and probes. Solid colored lines indicate the average perceived separation between probes. Transparent rectangles indicate the 95% confidence interval of the mean. The colored dots indicate individual participants. The dashed horizontal line at 4 dva indicates the frame movement amplitude. The “same plane” condition has no depth offset, and we can see that the three other conditions have a mean that falls in the 95% confidence interval of this control condition, indicating no effect of these depth offsets.

The lack of any decrease in effect with depth separation is in contrast with Experiment 1, where a decrease in the perceived separation was found with increasing horizontal and vertical offsets. There could have been a decrease in perceived probe offset here with even larger depth offsets but there appears to be no effect for the depth offset we used here.

## Effects in time

Does the frame effect extend in time before and after the presence of the moving frame? Here the various effects of time on the frame’s effect are evaluated in two experiments. First, the perceived offset between the probes is measured when the frame and probes are not on the screen simultaneously or overlap partially or overlap completely. Then, the perceived offset is measured for probes that are flashed at different time points within the frame’s left-right motion cycle, both in a frame in continuous motion and in a frame in apparent motion.

### Exp 3. Temporal offset

If the frame effect on position depends on grouping the probes with the frame, the frame may have to be present when the probes are flashed to see any effect. However, the grouping of the frame and probes and the spatial reference provided by the frame may persist for a brief time after the offset of the frame. Equally, the locations of probes that precede the frame may retroactively be referenced to the frame if it appears soon enough. Here, the probes are presented either while the frame is present or before or after it has appeared. The effect of the number of times the frame repeats its left-right motion cycle is also varied.

#### Methods

The frame cycled back and forth 1, 2 or 3 times and probe pairs were presented at three different times relative to the presentation of the frame. In the first, both probes appeared during the last displacement of the frame; in the second, the probe pair straddled the onset or offset of the frame’s presentation so that only one of the probes appeared while the frame was present; and in the third, there was no overlap between the probes and the frame, both flashed either just before or just after the frame’s presentation. In the complete overlap condition, both probes would be inside the first cycle of the frame (for the before/pre condition) or the last (for the after/post condition). When there was only one cycle of motion, the before and after conditions were the same. In each trial, the complete sequence of stimuli was repeated, with a gap of between 1.8 seconds and 3.15 seconds between repetitions, until participants finalized their response. The frame motion was always 4 dva per 250 ms. A demonstration movie, S1, is available in the Supplementary Materials (https://cavlab.net/Demos/FEAST/#4/). Combining the three levels of frame cycle repetitions with 2 orders (probe presentation of probes before and after), and 3 degrees of overlap (0, 1, and 2) created 18 conditions. Each of these conditions was shown twice during each of four blocks for a total of 8 repetitions to produce a total of 144 trials. In addition, 4 standard control trials with a single motion cycle were added to each block to test the effects of time-on-task with frame motion amplitudes of 0.8, 1.6, 2.4 and 3.2 dva but no temporal offsets (16 trials). Reports from these control stimuli are analyzed in Experiment 8.

#### Results and Discussion

The magnitude of the perceived separation between the two probes was similar in the standard condition here (frame present when both probes were flashed) compared to the previous two experiments. The offset settings (Figure 5) showed no notable effect of the number of repetitions, suggesting that there was no build-up of the frame-induced position shifts over repetitions of the motion cycles. Moreover, the frame-induced shift was only seen when there was some overlap (partial or complete) between the probes and the presence of the frame. This indicates that the effect arises only when the frame is present and its effect does not extend in time (no persistence or retroactive effect).

**Figure 5.**
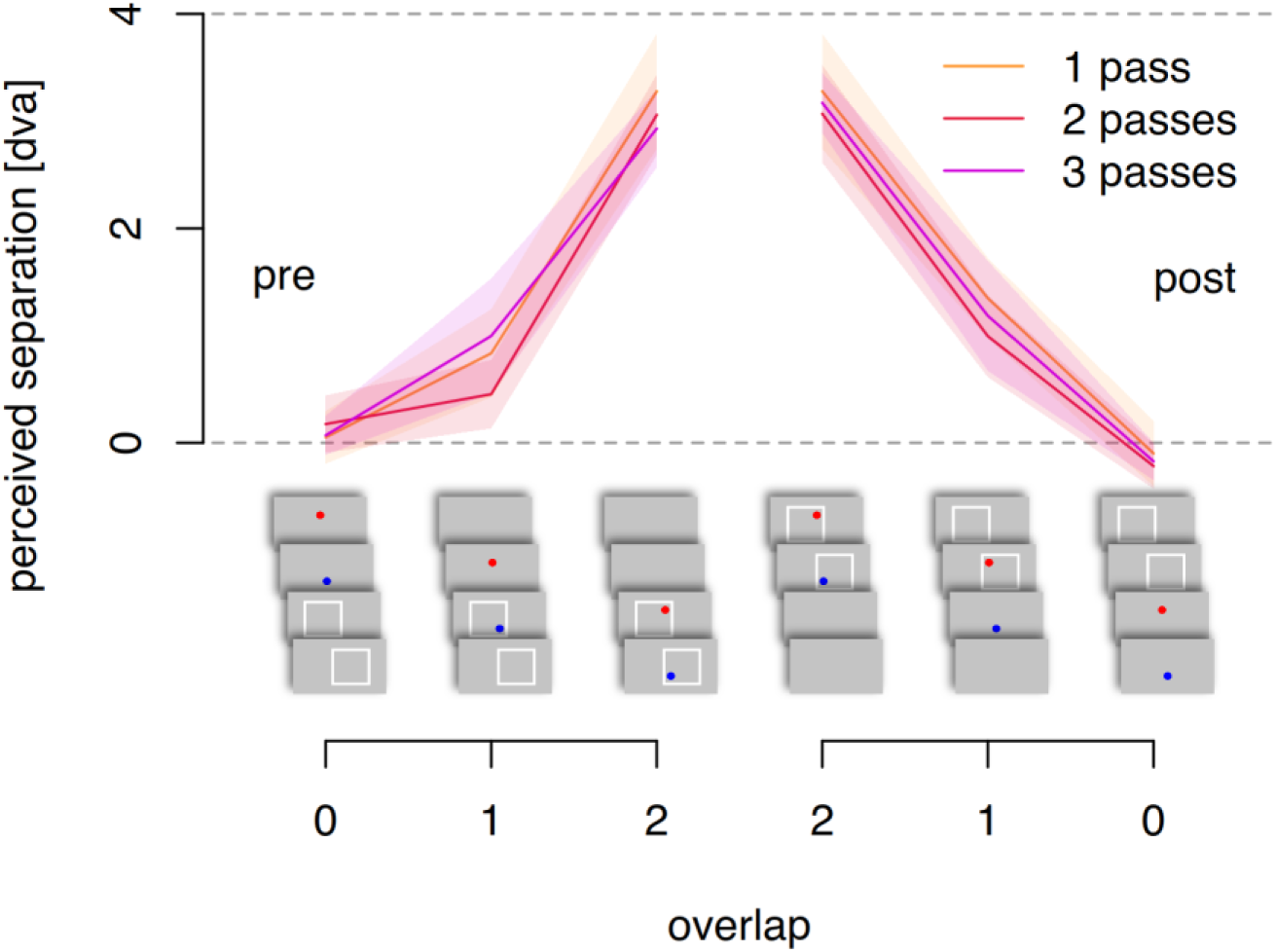
Perceived separation between the two probes as a function of the number of motion cycles and the overlap of the flashed probes with the presence of the frame (neither, one, or

A three-way, repeated-measures ANOVA confirmed that the number of frame passes did not have a main effect (F(1.24,16.12)=3.44, p=.075, η_G_^2^=.015), nor did it interact with any other factor (all p>.28). The temporal overlap between the frame and the probes did have a main effect (F(1.88,24.39)=310.67, p<.001, η_G_^2^=.813). Whether the probes were presented before or after the frame had no main effect but did interact with the temporal offset between the first/last frame pass and the time the probes were flashed (F(1.19,15.45)=5.98, p=0.023, η_G_^2^=.046). There were no other effects.

### Exp 4. Varying probe timing

Here we test the effect of delaying the probes’ appearance relative to the point of reversal of the frame’s motion with both continuous and apparent motion of the frame. Delaying the probe’s presentation changes its position relative to the frame, linearly reducing its offset from the frame’s center point. For example, when the two probes are flashed midway from the beginning of the cycle, they are both positioned in the middle of the frame and should have no frame-relative offset. Perceived offset should therefore decrease linearly with the delay of the probe’s appearance from the start of the motion cycle if the frame-relative position is the source of the perceived separation. The frame could either move continuously, as in the previous experiments, or be in apparent motion – presented only at its two end points and blanked between. To test the effect of delay for both continuous and apparent frame motion, we converted the delay between the motion reversal and the probes into probe offset in frame coordinates. Importantly, for the apparent motion case, the frame was not present between the endpoints and so the frame-relative separation is based on the interpolated position of the virtual frame, testing whether the frame has to be present for a frame effect to occur.

#### Methods

The frame motion was 4 dva and the motion duration was 333.3 ms with a pause at motion reversal of 133.3 ms. We used both a classic frame, showing the frame over its full path, as well as an apparent motion frame that was presented only at the extremes of the path (but appeared as a frame moving back and forth to participants). A demonstration movie, Movie S2, is available in the Supplementary Materials (https://cavlab.net/Demos/FEAST/#6/) Both types of frames were combined with 10 delays relative to the motion reversal: 4 leads, one synchronous, and 5 lags. These delays varied in steps of 33.3 ms, from –133.3 ms to +166.7 ms. Only the first 4 lags are included in the analysis, to balance the lead and lag data at each delay. Probes were flashed for 66.7 ms as in the other tasks. For the shortest leads and lags the probe flash overlapped for 50% with the stationary frame during the pause while the other half coincided with the frame motion, or the frame absence for the continuous and apparent motion cases, respectively. The 10 delay conditions and two motion types, were all repeated twice within each of 3 blocks for 120 main trials. These timings were used for only 8 of the 14 participants due to a coding error and only the data from these 8 are analyzed. In each trial the sequence of stimuli was presented continuously until participants finalized their response. In addition, the standard control trials were added to each of the 3 blocks to test the effects of time-on-task with frame motion amplitudes of 0.8, 1.6, 2.4 and 3.2 dva with the probe presentations synchronized with the motion reversals, and only in the continuous frame (2 repetitions of each in each block for 24 trials). Reports from these control stimuli are analyzed in Experiment 8.

#### Results and discussion

Figure 6 shows that the perceived offset between the probes decreases as a function of the horizontal distance between the probe positions within the frame (or interpolated frame in the apparent motion case). This is well described as a linear function for both types of frames (continuous: p<.001; apparent: p=.04). The data for each probe separation in frame coordinates are averaged over the corresponding lead and lag conditions as the probe separations are the same for lead or lag delays. The decrease is similar for both continuous and apparent motion cases although the overall size of the illusory separation is smaller for the apparent motion case, as has been previously reported (Cavanagh et al., 2022). The decrease for both types of motion is strongly linear with increasing delay and intersects the horizontal axis near the 0 dva probe separation in frame coordinates, as would be the case if the perceived offset were driven by the positions of the probes in the frame at the time they flashed.

**Figure 6.**
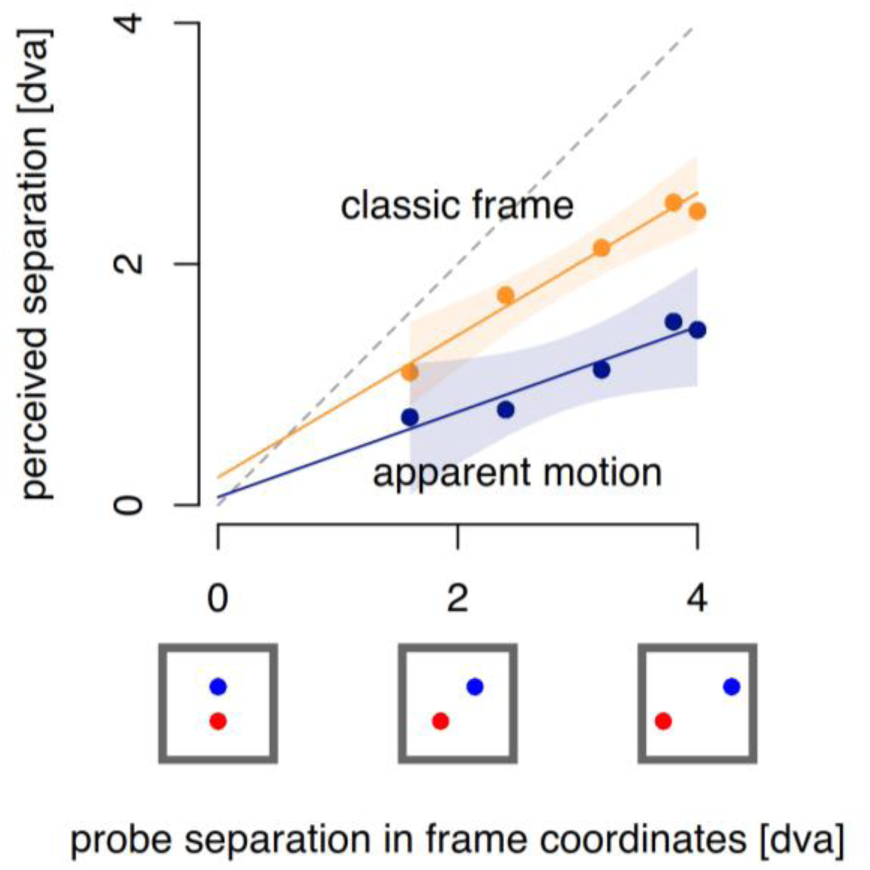
Perceived offset as a function of several probe distances in frame coordinates caused by delayed presentation relative to the motion reversal, averaged over leading and lagging delays.

The frame effect is also seen for the apparent motion condition where the probes are presented when the frame is not present. The effect could have disappeared for the delays when the frame was not present on the screen, but instead, we see a linear decrease, similar to that for the continuous frame motion where the frame was always present. Experiment 3 showed that there is no frame effect if the frame is absent when the probes are flashed but here the effect remains. This suggests that it is not the physical presence of the frame when the probes appear that is critical but the presence of an interpolated percept of a frame as is produced by apparent motion (Kolers, 1972). In Experiment 3, when the probes both precede or both follow the frame’s appearance, there is no impression of the frame when the probes appear. In contrast, with the apparent motion stimulus, the frames are absent between the presentation at the two end positions, but the impression of a frame is present.

## Effects of frame motion

Here we examine whether the critical factor in the frame’s effect is the motion of the frame’s outer boundary or the motion of the texture it encompasses. These two factors are the same in the case of the frames of the previous experiments which have no moving features other than the outer contour. To compare the outer boundary to inner feature motion, the frame was a square region of random texture where the texture could move with the boundary of the square region or remain static or move against it. In the first control experiment, the effect of an unbounded texture motion was compared to that of a frame with a single outer boundary across a six-fold range of speeds. In the second experiment, the frame was a square region of texture where the texture either moved with the frame or was static on the screen (the square region moved like a window revealing the static dots behind it), or the texture moved one way while the square area where the texture was present moved the other way. The following two experiments both consisted of a preliminary check of perceived extent of the motion path, followed by an assessment of the perceived offset between flashed probes. The motion and offset judgments of both experiments were run together but described and analyzed separately here.

## Effects of motion

### Exp 5. Frame and background motion

First, we tested if the grouping between flashed probes and frame’s motion requires the frame to have a bounded shape, with edges, or whether it can be a pattern or texture extending beyond the boundaries of the screen. MacLeod et al. (2024) had already shown that a moving random texture produces a perceived offset of two physically aligned probes that are flashed at the motion’s reversal. They found that there was an effect but smaller in magnitude than the classic frame effect that had been reported in an earlier paper (Özkan et al., 2021). We repeat this test and directly compare it to the classic frame effect under comparable conditions. We also asked participants to report how far the frame or texture appeared to move in the frame and texture conditions to check that the motion impressions were comparable and that the reduced effect reported by MacLeod et al. (2024) was not due to a reduction in the perceived extent of the motion path. Finally, we tested the effect of a fixation on the perceived separation. Up to now, all the experiments allowed the participants to move their gaze at will. Here, we also compared free viewing to fixation for the classic frame condition only.

#### Methods

The classic frame was the same as in previous tasks, a 7x7 dva square, with a 0.5 dva white outline (and a 6x6 dva blank center). The dots in the texture were 0.4 dva squares, with a lifetime of 1 s and with 30%, 40%, 60% and 70% gray level (of the monitor’s luminance range). This strip extended beyond the edges of the screen on the left and right, but was 6.8 dva high, covering almost the same vertical extent as the classic frame. The motion was 333.3 ms for each transit (excluding the 150 ms pauses in movement at the extreme positions).

We tested the perceived travel of the frame and the texture field seen for six physical displacements of 1, 2, 3, 5 and 6 dva all with 333.3 ms duration of motion and at 4 dva for 5 durations (200, 250, 333, 500 and 1000 ms). There was a 150 ms pause in movement at each motion reversal. There were no probes flashed during these path length trials. Participants used the mouse to adjust the length of a horizontal bar on the top right to indicate how far the stimulus appeared to have moved from one end to the other of its path and pressed the space bar to finalize their answer. There were 40 trials for the length judgments (1 repetition for 2 stimuli and 5 path lengths at 333 ms duration and 3 repetitions for 2 stimuli and 5 durations at 4 dva path length) in each of the 3 blocks.

**Movie 3.**
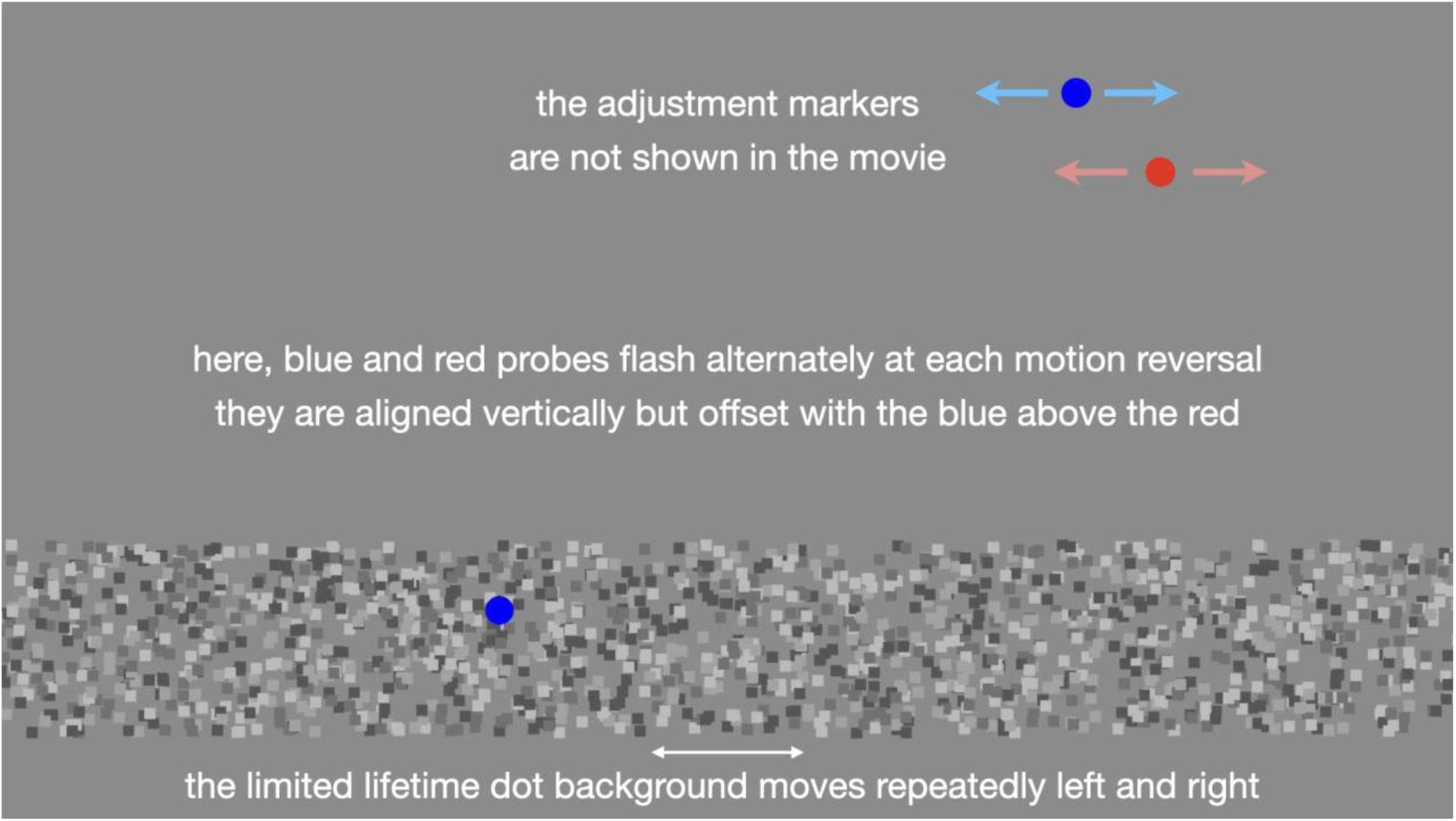
Classic frame motion with and without fixation versus extended limited lifetime, random dot motion. The adjustable dots in the top right corner are not shown in this demonstration movie but were present in the experiment. Use this link to open the movie in a browser window: https://cavlab.net/Demos/FEAST/#8/.

The main task conditions used the same stimuli with a single, fixed 4 dva distance of travel for the frame and texture field, but now with the probes flashed at the motion reversals. Participants indicated the perceived separation between the two probes by adjusting the slant between two reference dots as in the previous tasks. The probe separation trials were presented at one of five different movement durations (200, 250, 333, 500 and 1000 ms). Each of the 10 conditions (2 stimulus types and 5 durations) was repeated 2 times in each of the 3 blocks plus 4 control trials, making a total of 72 trials.

We also examined any difference between peripheral viewing (with a fixation point) and free viewing (with occasional foveation of the stimulus). We used free viewing in all other tasks, but here we included stimuli with the classic frame where we asked participants to fixate a dot at the center of the screen (in between the stimuli and the reference probes) when judging the slant between the dots. Participants were asked to fixate this dot – if it was present - while observing the stimuli. They were told that they could look at the reference probes while adjusting them, but they should not look directly at the frame and probes stimulus. If the fixation dot was not present, they could look directly at the stimulus. They could look at the reference probes in all conditions.

This task also included the standard control stimuli at different amplitudes (0.8, 1.6, 2.4 and 3.2 dva) in the standard frame to evaluate the effect of prolonged experience with the illusion (see Experiment 8).

#### Results and discussion

The reported motion path length is shown in Fig. 7A and scales roughly linearly with the physical motion amplitude for both stimuli (F(3.23, 42.03)=69.94, p<.001, η_G_^2^=.521) with the perceived path length slightly less for the texture than the frame (F(1, 13)=7.93, p=.015, η_G_^2^=.049), with no interaction. So, although reduced, the moving random texture field produced a strong impression of displacement. The perceived path length is longer than the physical length, a finding at odds with the previous papers reporting path shortening for stimuli in motion (e.g., Sinico et al., 2009; Cavanagh & Anstis, 2013). It is possible that the lengthening seen here is a result of the difference in size judgments between the fovea and the periphery where tests in the periphery appear smaller than in the fovea (e. g., Newsome, 1972; Baldwin et al., 2016). In the case of the experiment here, whenever the participant was looking at the test stimuli in the moving frame, the matching bar would be in the periphery and the apparent length would be smaller, leading to an adjustment longer than the veridical length to make a match.

**Figure 7.**
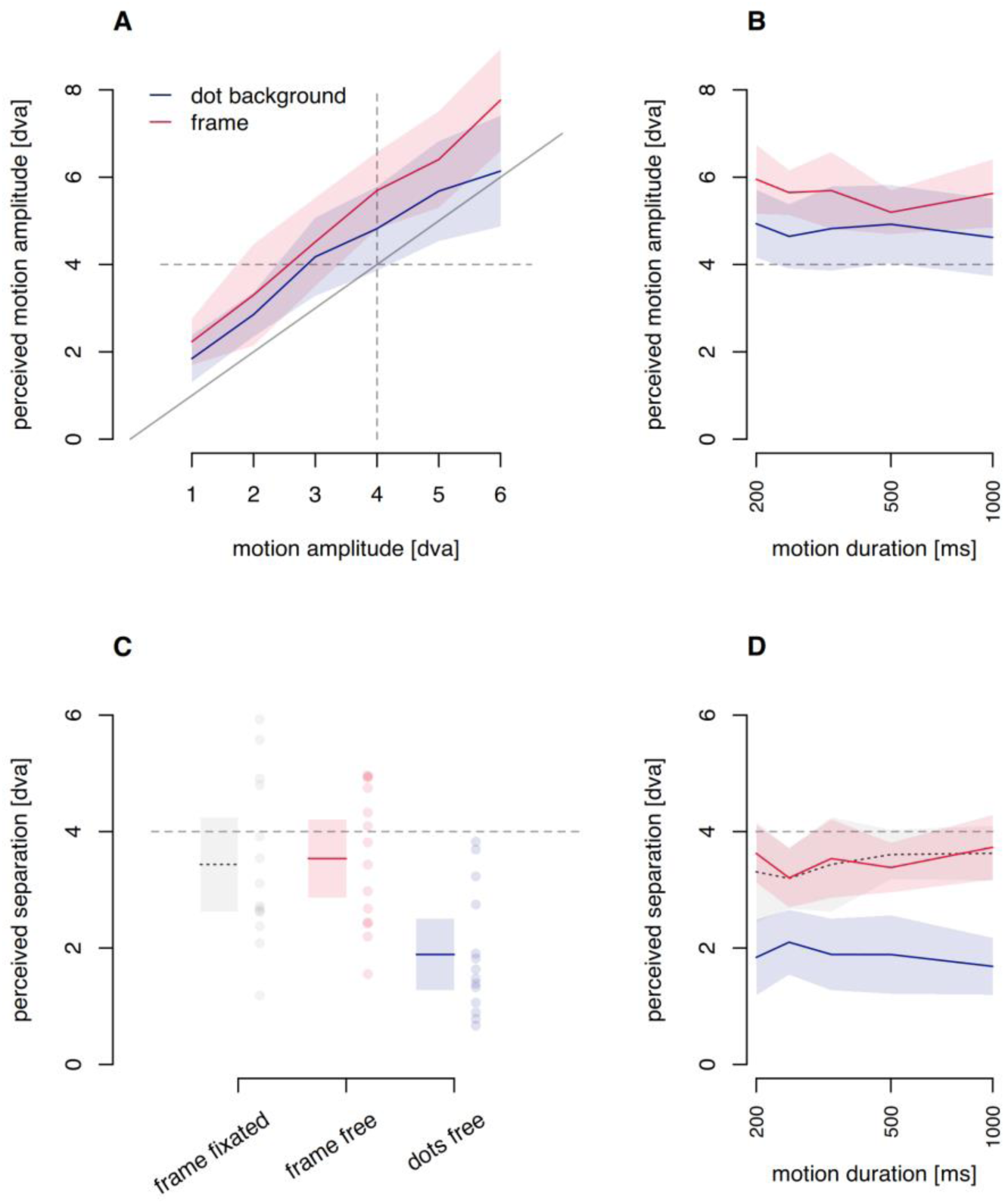
A. Perceived path length for the classic frame (in red) and for the full texture field (in blue) as a function of physical motion amplitude. The diagonal line indicates veridical judgments. B. Effect of motion duration on perceived motion amplitude. C. Perceived probe separation for the classic frame during fixation (gray), for the classic frame with free viewing (red) and for the dot field with free viewing (blue) (frame fixated; gray). D. Perceived probe separation as a function of motion duration. Colored lines in all panels indicate averages in the conditions and the shaded areas denote 95% confidence intervals. In panel C, the dots indicate individual participants.

The classic frame created a perceived motion amplitude of 5.6 ± 1.0 dva (Fig 7B, red line), while the moving, frameless texture (Fig. 7B, blue line) created a perceived motion amplitude of 4.8 ± 1.3 dva across the 5 different movement times. Both percepts are different from 0 (p<10^-8^) and the motion induced by the frameless texture motion is smaller than that induced by the classic moving frame (t(13)=3.47, p=.004).

After characterizing the motion percepts these stimuli induce, we now test their effect on the perceived offset between flashed probes (Fig. 7C&D). While there is some perceived separation between the probes in the random dot background of 1.9 dva (t(13)=6.63, p<.001) this is much smaller than the 3.5 dva separation seen with the classic frame in free viewing (t(13)=5.65, p<.001). These results replicate those of MacLeod et al. (2024).

As for the effect of fixation with the classic frame (Fig. 7C), there was no significant difference (t(13)=0.28617, p=0.779). A Bayesian t-test (with a non-informative prior) shows that there is moderate evidence that the illusion is equally strong in both conditions (BF_10_=0.28). This means that the frame effect is similar with peripheral viewing and free viewing (with occasional foveation).

The perceived offset follows the same pattern across all motion durations (Fig. 7D).This is confirmed with a 3x5 repeated measures ANOVA with stimulus type (free viewing frame, frame with fixation, dot background) and motion duration (200, 250, 333.3, 500 and 1000 ms). There is an effect of stimulus type (F(1.98,25.78)=53.88, p<.001, η_G_^2^=.361). There is no effect of motion duration and no interaction with stimulus type.

This experiment shows that while there is a small decrease in perceived motion in the dot background, it also leads to a much larger decrease in perceived offset between the flashed probes. This indicates that a moving, bounded frame rather than a moving, extended background leads to stronger frame effects.

### Exp 6. Internal frame motion

In the previous experiment, we found that the texture’s motion on its own does produce some probe separation. To test if this can modulate the frame effect, we now place a texture inside a bounded frame and have the texture motion and the outer boundary motion compete in their influence on the perceived probe separation. Here we construct a frame that is a 6.8 dva square region of random dot texture. The square region moves left and right as in previous experiments while the texture within the square can move with the square, against it, with it but at twice its speed, or remain static on the screen. These variations of the frame may affect the perceived separation of the flashed probes differently but may also affect the perceived travel of the frames, and consequently the position shifts as well. To control for this, participants also reported the perceived distance traveled by the frame.

#### Methods

The square region filled with dots covered a 6.8x6.8 dva area. The dots were 0.4 dva squares, with a lifetime of 1 s and had randomly distributed luminance levels of 30%, 40%, 60% and 70% gray (in the monitor’s luminance range). The “frame” moved through 4 dva in 333 ms and this was combined with four kinds of internal motion. First, dots could move with the frame. Second, dots could be stationary on the screen, i.e. not moving with the frame. Third, the dots could move in the opposite direction of the frame. Fourth, the dots could move in the same direction as the frame but twice as fast. In stimuli where the dots did not move with the frame, dots that exited the frame region were randomly repositioned within it as were dots after their lifetime expired. The apparent distance the frame traveled was also tested separately from the probe separation they evoked using the method of the previous experiment.

**Movie 4.**
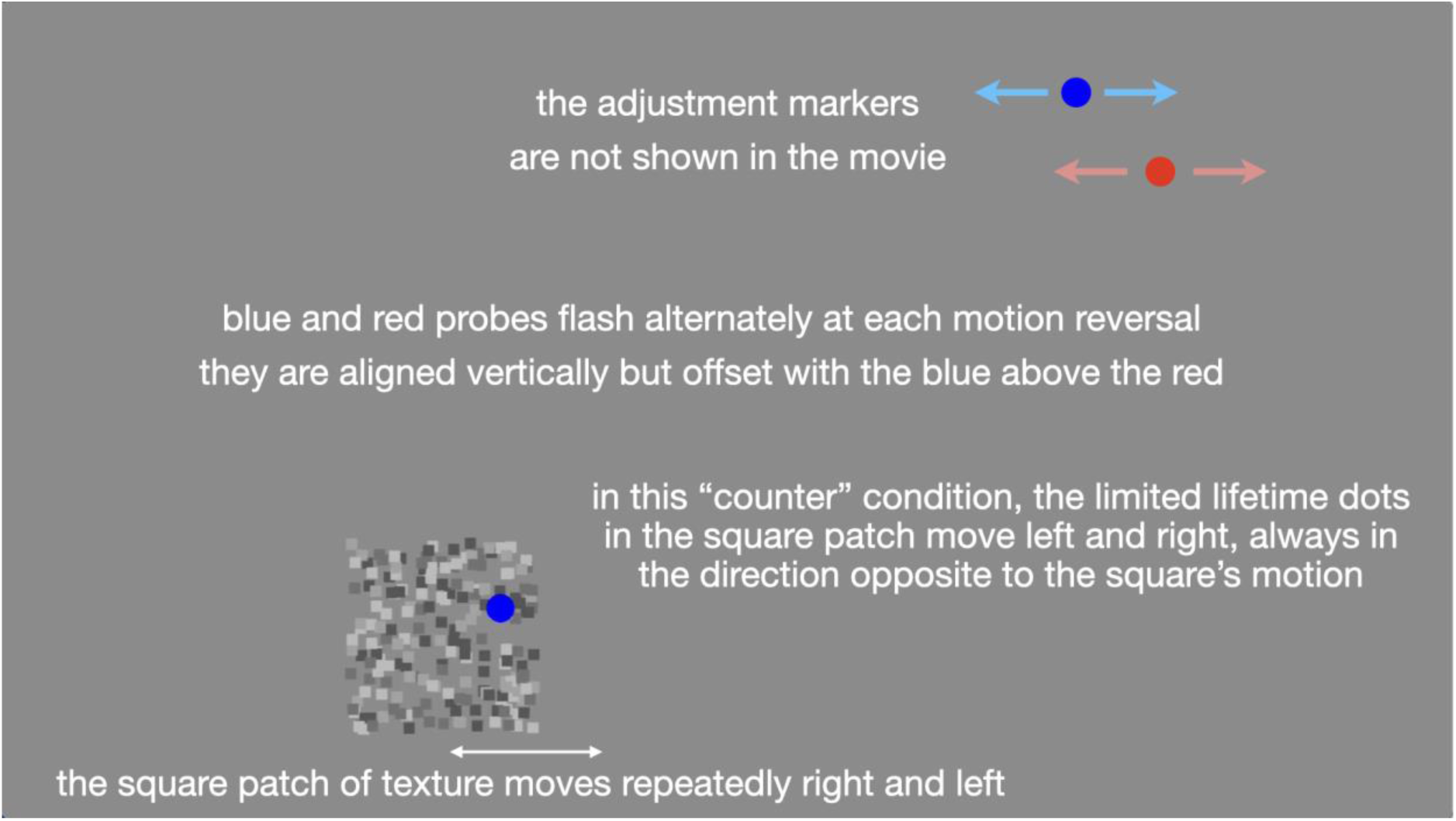
Frames made of random dots. The internal texture could move against the textured region’s motion, remained in place, moved with the region’s motion at the same speed or at twice the speed. The adjustable dots in the top right corner are not shown in this demonstration movie but were present in the experiment. Use this link to open the movie in a browser window: https://cavlab.net/Demos/FEAST/#10/.

In the first task, participants judged the apparent amplitude of the frame’s motion path without probes present, as in the previous experiment. This task had 3 blocks where each of the 4 stimuli were shown 3 times per block for 36 trials.

In the second task, the probes were presented, and participants judged the perceived offset between the two probes as before. This task had 3 blocks where each of the 4 stimuli were shown 4 times per block for 48 trials.

The control trials to test the frame effect across time-on-task were present during this second task. The classic frame stimulus had movement amplitudes of 0.8, 1.6, 2.4 and 3.2 dva and a motion duration of 333 ms with two trials per block, for an additional 4 trials per block, for a total of 12 control trials.

#### Results and discussion

When the dot field is present within a moving region (Fig. 8A), the apparent distance traveled by the region was unaffected by the relative motion of the dots (against, static, with, with the region at double speed, F(1.70,22.07)=1.86, p=.184, η_G_^2^=.044). This indicates that perceived frame travel is relatively matched across conditions and would not contribute to any difference in the perceived separation of the two flashed probes.

**Figure 8.**
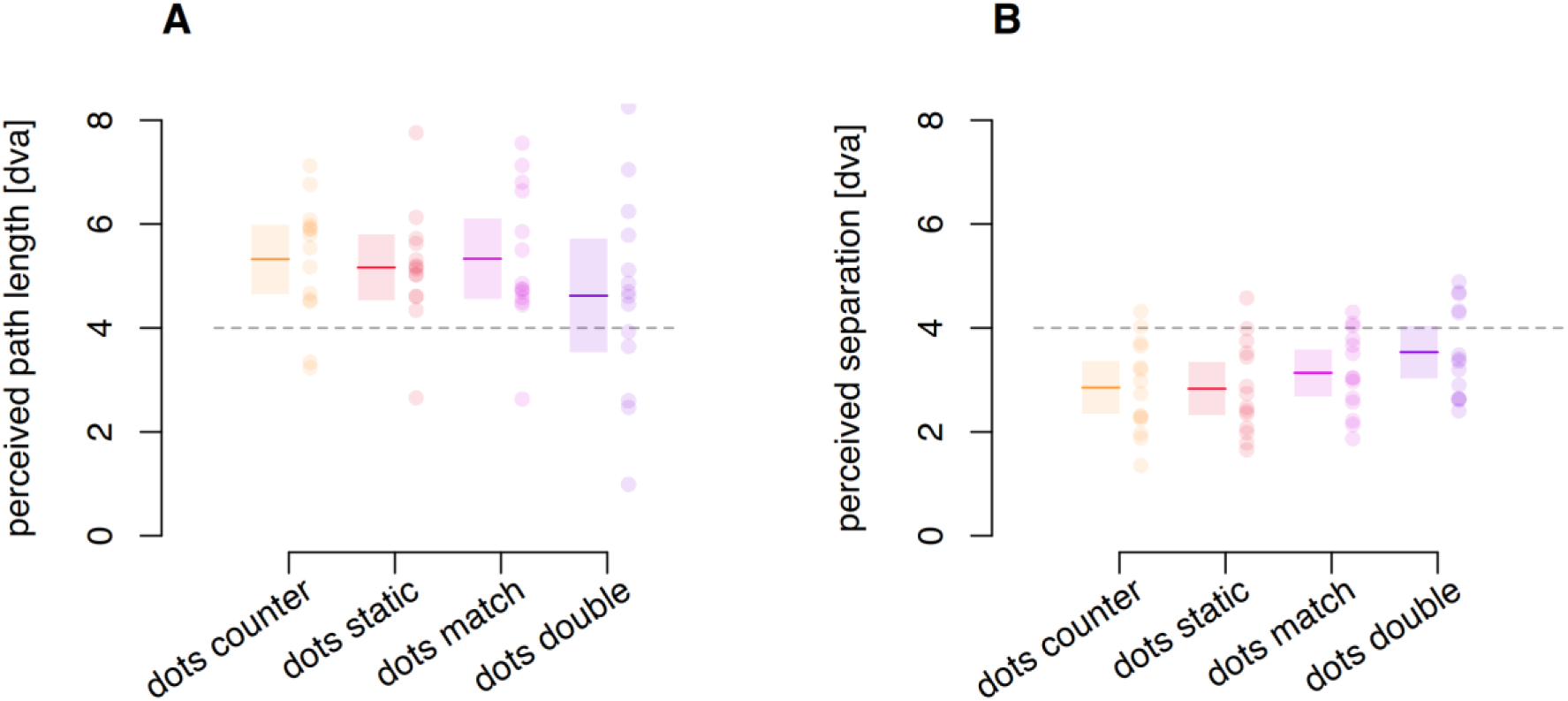
A. Reported length of motion travel for the four types of frames consisting of limited lifetime dots. Dots counter: motion of dots is opposite to that of the frame. Dots static: no internal motion relative to the screen. Dots match: dot motion matches the frame motion. Dots double: dot motion is double that of the frame and in the same direction. B. Perceived probe separation induced for each of the four types of frames. Horizontal dotted lines indicate the physical frame travel of 4 dva. Colored lines indicate averages within each condition. Shaded areas denote a 95% confidence interval for each condition. Dots represented individual participants.

The effect of the frames on the perceived dot separation appears similar across the four frame types (Fig. 8B) but did differ significantly (F(2.68,34.86)=6.65, p=.002, η_G_^2^=.105). Post-hoc contrasts (Sidak adjustment for multiple comparisons) showed that the frames with the double-speed internal motion generated a greater perceived separation (mean of 3.53 dva) than the static dots (2.83 dva, t(13)=4.31, p=.005) or counter moving dots (2.85 dva, t(13)=4.12, p=.007). No other contrasts were significant (all *p*>0.16).

The results of Exp. 5 indicate that background motion by itself can create a shift in perceived location, albeit weaker than the effect of an actual frame. The current experiment (Exp. 6) shows a modulation of the frame effect by internal motion that is much smaller than the effect of the texture motion on its own. Combined, the results of Exp. 6 and Exp. 5 indicate that the displacement of a bounded region dominates in generating the perceived probe separations, even in the presence of incongruent internal motion signals. While the local motion signals on their own do produce separation of the probes, their contribution seems to be largely vetoed in the presence of the bounded frame. That is, the frame effect depends predominantly on the displacement of the frame itself, not on its motion signals.

### Exp 7. Frames moved by the participant

Here, the frame’s motion was yoked to the hand motion of the participant (using an auditory metronome-like guide for timing) and the strength of the frame effect was compared for hand-generated motion against that seen with standard frame motion.

#### Methods

The (7x7 dva) frame moved in three different conditions: 1) a stylus controlled by the participant moved the frame, which moved in the same direction as the stylus; 2) the frame was again controlled by the stylus but the motion was reversed so a rightward stylus motion produced a leftward frame motion and vice versa; and 3) the frame was under computer control and the participants were instructed to keep their hand still to prevent any efference copy. In all three conditions, a metronome clicked every 500 ms to indicate the timing for the motion reversals. A probe was briefly flashed at each motion reversal, aligned horizontally but offset vertically as in previous experiments. To control the frame motion, participants moved a stylus with their right hand back and forth between the two ends of a slot attached to a drawing tablet, reaching the end of the travel and reversing motion in step with the metronome. A full sweep of the slot with the stylus produced 4 dva of frame travel. The computer-controlled frame also traveled 4 dva each 500 ms. Participants reported their percept by pressing the left and right arrow keys on a keyboard with their left hand to adjust the standard reference dots to match their percept of the flashed probe locations. They finalized their setting by pressing the up-arrow key. Each of these conditions was repeated 5 times per block, such that each of the 3 blocks consisted of 15 trials, and each condition was measured 15 times for a total of 45 trials. There were no control stimuli here to test the effect of time-on-task.

#### Results and discussion

The results (Fig. 9) show the perceived probe separation for the three types of frame motion: congruent with hand motion, computer controlled, and opposite to the hand motion. There are no significant differences between the conditions (F(1.45,18.85)=1.62, p=.224, η_G_^2^=.009). The additional information from the efference signals of the hand motion do not affect this percept: the frame effect is a purely visual illusion.

**Figure 9.**
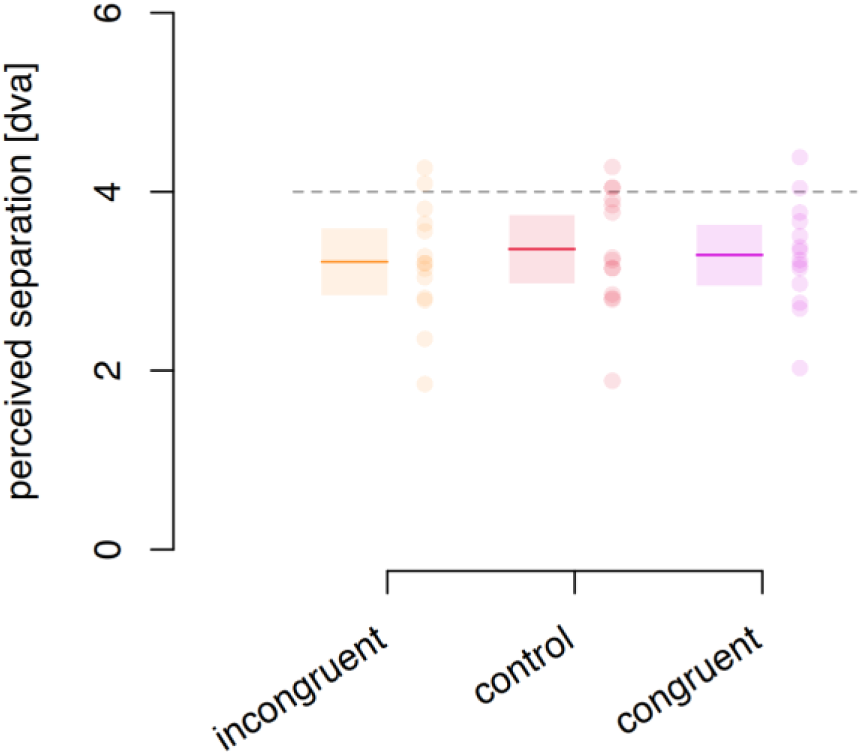
Effects of self-moved frames. Participants reported perceived separation of the probes that were moved in the direction opposite their hand motion (incongruent), moved by the computer (control), or moved with their hand motion (congruent). The dashed gray line indicates the actual extent of the frame motion. Colored lines indicate the averages within conditions. Shaded areas denote the 95% confidence interval of the mean. Dots represent individual participants.

**Movie 5.**
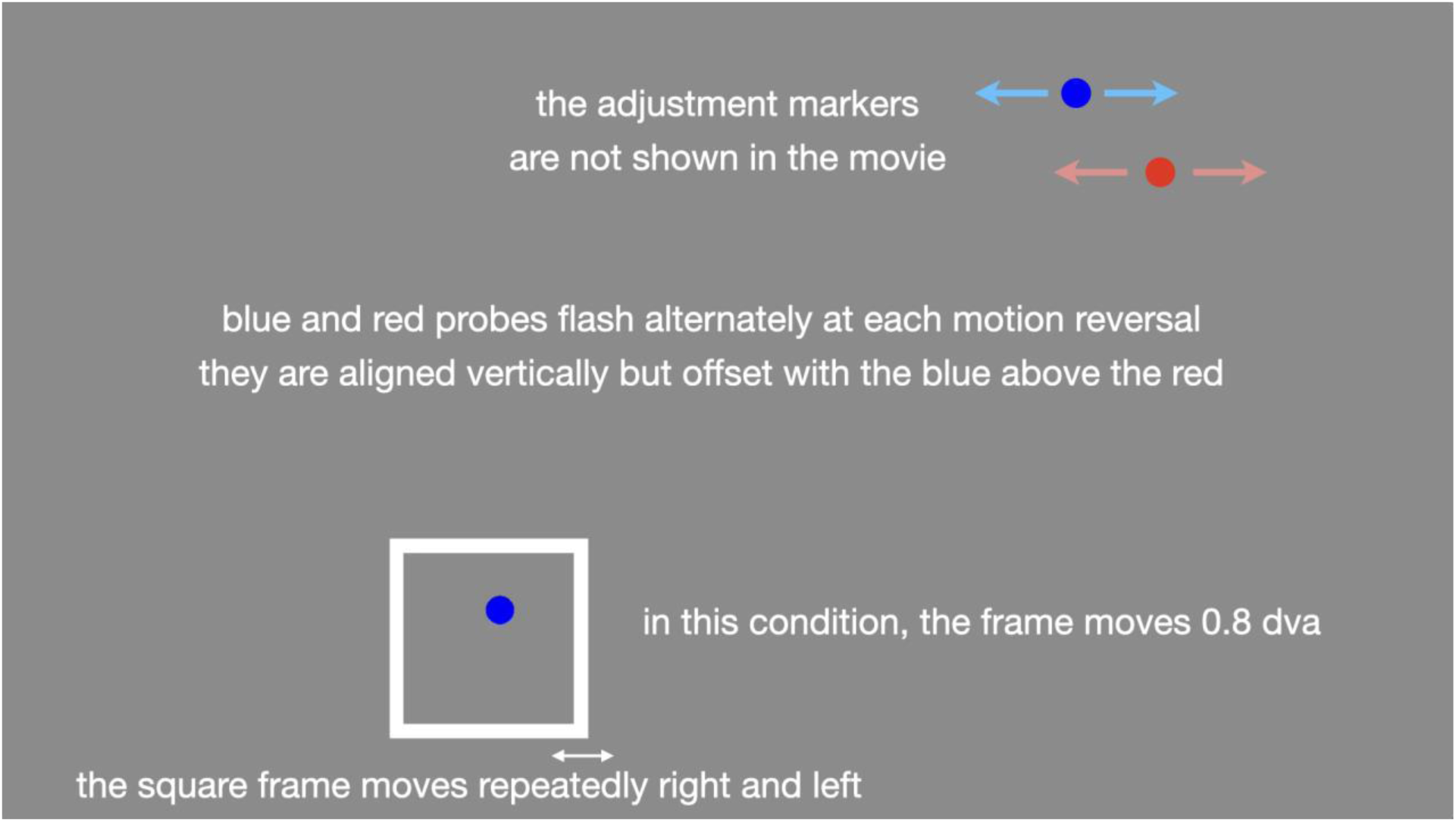
Control measure of the strength of the frame effect across time. The perceived probe separation was measured for 5 path lengths of the motion. These trials were interspersed with the main experiment trials across Experiments 1, 3, 4, 5, and 6. The adjustable dots in the top right corner are not shown in this demonstration movie but were present in the experiment. Use this link to open the movie in a browser window: https://cavlab.net/Demos/FEAST/#12/.

## Time-on-task

### Exp 8. Time-on-task and illusion strength

Does the frame-induced position shift decrease with continued testing over time? All participants performed the 7 experiments in a single session spanning an average of 2 hours but in different orders for different participants (range: 80 minutes to almost 3 hours). As many breaks as wanted could be taken in between tasks and blocks, and breaks could be as long as wanted. The strength of the frame effect was monitored across the duration of testing by inserting trials of a control frame task in Experiments 1, 3, 4, 5, and 6. The timing of the control tasks was evaluated based on the start times of individual trials relative to the start of each participant’s session.

#### Methods

The classic frame stimulus was presented with five different motion amplitudes (0.8, 1.6, 2.4, 3.2, and 4 dva). The shorter path lengths, 0.8 to 3.2 dva, were the control trials added to the experiments. The data for path length of 4.0 dva were taken from the main experiments for the conditions that matched the control trials. For all of these trials, the frame was 7x7 dva, with no spatial, temporal or depth offsets and probes flashed at the motion reversals. The frame motion cycled continuously until the participant responded except in Experiment 3 where the frame made a single return cycle of motion (left then right or right then left, randomly). and these single cycles were repeated with a gap of 3.15 seconds between them until the participant responded. The duration of frame motion was 250 ms in Experiments 1, and 3, but 333 ms in Experiments 4, 5, and 6. This change in motion duration affected the frame’s speed. Previous results (Özkan et al., 2021), and Exp 5 here, showed that the frame’s effect is independent of speed so the data with different speeds here are not analyzed separately. In Experiment 4, the first 7 participants used a motion amplitude of 1.8 dva instead of 1.6 dva due to a technical error, and these 78 trials are not included in the analysis of a total of 1381 trials. In each condition within each epoch, whenever a participant had more than one data point, these multiple values were replaced with the median of these repetitions. A total of 201 values remained following this data compression step.

#### Analysis

The responses were binned into four sets according to the time of the response relative to the start of the first task for each participant. Each bin represented a half hour of experiment time, allowing us to track the responses over elapsed time. The control trials are distributed randomly across the full duration of each of the 5 experiments that had embedded controls. Since only a few time-on-task stimuli are included, many participants do not have reports for each motion amplitude in each half-hour period. To analyze these data with many missing values, we fitted a linear mixed effects model with perceived offset as the predicted variable, with the 4 time epochs and 5 frame motion amplitudes as fixed effects and with participant as a random effect. For interpretability, this is converted to ANOVA-like output using the Satterthwaite method (Luke, 2017; Satterthwaite, 1941).

#### Results and discussion

Control trials were inserted in 5 of the experiments of each participant’s session. These presented the classic frame stimulus and varied the amplitude of the frame’s motion. Figure 10 plots the average perceived separation reported for these 5 motion amplitudes across the four successive, 30-minute epochs.

**Figure 10.**
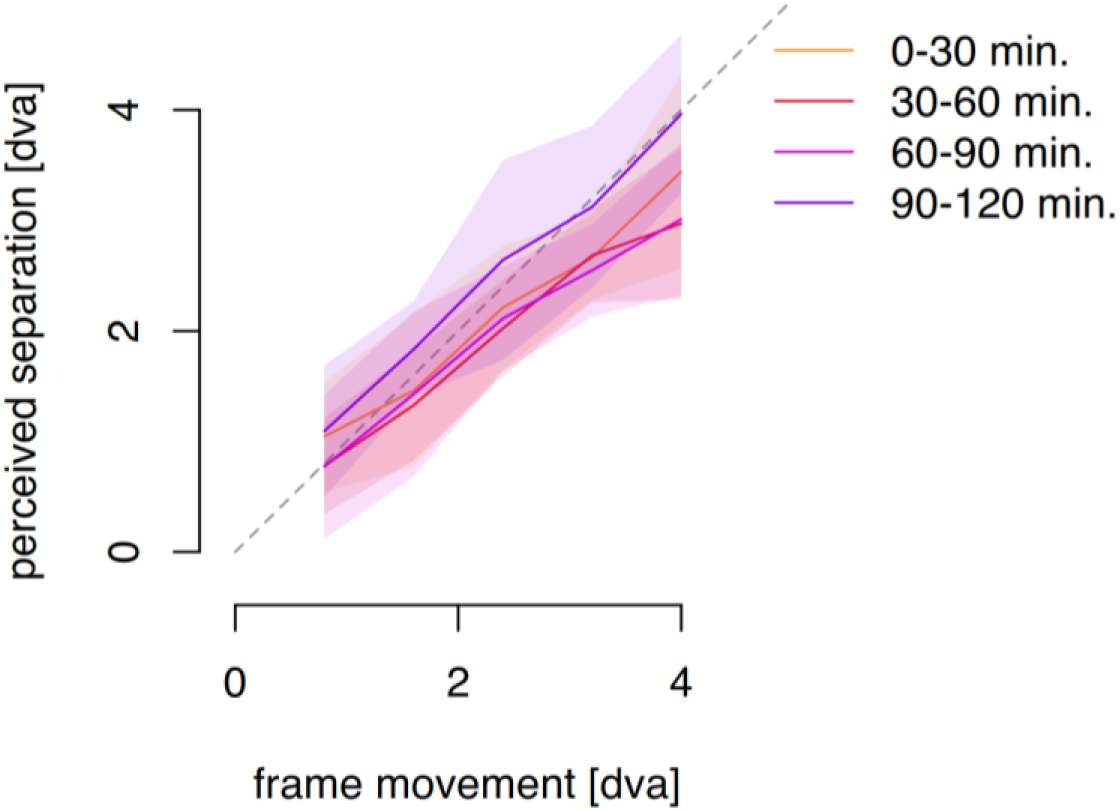
Perceived offset in the control classic frame effect across the 5 motion amplitudes, shown separately for each of the successive 30-minute epochs. The average across available data within each half hour period is represented by a solid, colored line. The shaded areas denote 95% confidence intervals of the mean. The dashed gray line indicates unity between the actual frame motion amplitude and the reported percepts (100% illusion).

In Figure 10 we do not see any modulation of perceived separation across the 4 time epochs, and the Linear Mixed Effects model with Satterthwaite approximation confirms this impression. There is the expected effect of the frame’s motion amplitude on perceived separation (F(1,183.99)=41.85, p<.001) which reached almost 100% of the frame’s travel across conditions and motion amplitudes. There is no effect of time epoch (F(1,84.64)=0.274, p=.601) and no interaction (F(1,184.00)=0.688, p=.408). Continued exposure to the frame stimulus does not affect the amplitude of the perceived shift.

## General Discussion

Overall, we found that the influence of the moving frame on the perceived locations of flashed probes did extend in space but not in time. Specifically in space, the separation seen between two probes was large when the probes were within the frame but dropped progressively as the probes moved outside the frame, whether in the direction of the frame’s motion or orthogonal to it (Exp 1). The effect was lost once the probes were 9 dva from the center of the frame’s path. The frame effect also held when the probes and the frame were in different depth planes (Exp.2). There appears to be a region of influence around a moving frame that is not limited to the space enclosed by the frame.

In terms of the temporal extent of the moving frame’s influence, the result was completely different. There were no frame effects when the probes both appeared before or after the frame (Exp. 3). There was a small effect when one probe appeared when the frame was present while the second preceded or followed the frame (the probe pair straddled the onset or offset of the frame’s motion). There was therefore no evidence of a persistence or a retroactive effect for the influence of the frame’s motion. Note that in the case of apparent motion (Exp. 4), the probes appear in between the frame’s appearances at the two ends of the motion path, with no frame present and still produce a strong frame effect. Although the frame is not present when the probes appear, the impression of motion between the two end points of the frame’s positions is in place when the probes appear. This means that the position of the probes in the frame (seen or interpolated) determines the perceived offset between them. We did not analyze the difference between effects of lags and leads on the perceived separation. There could be differences due to other effects such as the flash lag or flash grab illusions (Nijhawan, 1994; Cavanagh & Anstis, 2013) that would vary based on the direction of motion. This will be examined further in future research.

Experiments 5 and 6 showed that the effect of the frame was not due to low-level motion signals but to the displacement of the frame. The motion of a background texture without a moving border did cause a perceived separation between the two probes, but smaller in magnitude than the classic effect of the square frame, replicating the results of MacLeod et al. (2024). However, when a square region of texture moved back and forth, it generated a large separation between the probes relatively independently of the motion of the texture within it. While there was a small modulation by the internal motion, the displacement of the frame’s border dominated over any effects from the motion of the texture within it. Along with the preserved effect in the apparent motion frame, this suggests that the effect relies primarily on the displacements of the bounding borders of the frame, and only to some extent on motion signals.

That is, in the experiments conducted here, the perceived offset between the probes was determined by their position within the frame at the time the probes were flashed – not by the frame’s motion signals. This is somewhat at odds with the previous finding that a complex frame motion path can reduce the frame effect (Cavanagh et al., 2022). It may be that position of the probes within the frames is less clear if the frame moves irregularly, or it could be that irregular motion of an object discounts it as a reference for other objects to some degree. This remains to be tested. Although we argue that the perceived positions are set by the locations relative to the frame, effects of apparent motion between the two probes may be contributing. The subjective experience of most observers is of two separate points without apparent motion between them but some do report the probe jumping back and forth. Here we have focused on the apparent offset between the two flashes without asking whether the participants experienced any motion. We have assumed that if there is any apparent motion, it is a consequence of the illusory offset positions, not the cause of them. A strong argument against apparent motion being the cause rather than the effect is that the illusion is still equally strong with frame movements taking over 1 second (Özkan et al., 2021, Fig. 5A, slowest speed). This temporal offset is beyond the ISIs at which apparent motion can be perceived (Kolers, 1972). These observations argue against apparent motion of the probes playing a role in creating the illusion.

Experiment 7 showed that the frame effect depended on the physical motion of the frame independently of whether its motion was yoked to the motions of the participant’s hand, or set by the computer, or moved opposite to the participant’s hand motions. There was no evidence for an effect of self-initiated motion seen in some other visual perception experiments (Beets et al., 2010; Dewey & Carr, 2013; Wohlschläger, 2000; Maruya et al., 2007; Veto et al., 2018; Skora et al., 2021). As a result, the frame effect seems to rely solely on the visual input.

Finally, there was no decrease in the size of the illusory separation over 2 hours of experimentation (Experiment 8). Other illusions decrease with extended testing (e.g., Porac, 1989) indicating that they are considered errors by the visual system and can be minimized with continued exposure. The frame effect is maintained over time and does not adapt away with continued exposure. This suggests that the effects of moving frames are something regularly encountered by the visual system and that the shifts produced by the frames are part of the visual system’s strategy for dealing with multiple frames of reference in the visual scene.

## Conclusions

When probes are flashed in a moving frame, their locations are seen in the frame’s coordinates – for example, red on the right, and blue on the left of the frame in Figure 1 – as if the frame were present and stationary at the center of its path (Özkan, et al., 2021). Surprisingly, this leads to very large illusory shifts, as much as 100% of the distance the frame travels. The experiments here revealed that the frame’s effect extended outside the contours of the frame by several degrees of visual angle, both to the sides of the frame in the direction of the motion and equally well above and below it. It was also undiminished when the probes and the frame were in different depth planes. Unlike the extension in space, the influence of the frame’s motion showed no extension in time – no persistence of the effect once the frame was removed and no retroactive effect before the frame appeared either. The frame effect was driven primarily by the displacement of an object, the frame, not its motion signals. The effect was strongest for moving bounded frames compared to moving unbounded texture. When the bounded region had an internal texture that could move with or against the frame’s motion or remained static, the displacement of the frame dominated and the internal motion was mostly ignored. The effect was unaffected by whether the motion was generated by the participant’s hand motions or not and was not reduced in strength across 2 hours of testing.

We show that the frame effect may be a consequence of the way the visual system deals with moving reference frames. If this is the case, the requirement for frames to be bounded and to rely on the position of the frame when probes are flashed may also reveal how the visual system partitions a dynamic scene to estimate the location of its parts. The effect does not rely on efference copies or expected motion, depending instead on purely visual and simultaneously presented stimuli.

## Supporting information

Movie 1 - stimulus + trial

Movie 2 - frame offsets

Movie 3 - dot backgrounds

Movie 4 - dot frames

Movie 5 - frame motion amplitude

Movie S1 - pre-post probes

Movie S2 - lead-lag

## Acknowledgments

This research was supported in part by grants from the Natural Sciences and Engineering Research Council of Canada grant RGPIN-2019-03938 (PC).

## References

Anstis, S., Cavanagh, P. (2024). Influence of frame and probe paths on the frame effect. Journal of Vision, 24(11). doi: 10.1167/jov.24.7.11.

Asch, S. E., & Witkin, H. A. (1948). Studies in space orientation: I. Perception of the upright with displaced visual fields. Journal of Experimental Psychology, 38, 325–337. doi: 10.1037/h0057855

Baldwin, J., Burleigh, A., Pepperell, R., & Ruta, N. (2016). The perceived size and shape of objects in peripheral vision. i-Perception, 7(4), 2041669516661900.

Beets, I. A., ’t Hart, B. M., Rösler, F., Henriques, D. Y., Einhäuser, W., & Fiehler, K. Online action-to-perception transfer: only percept-dependent action affects perception. Vision Research, 50 (24), 2633–41, doi: 10.1016/j.visres.2010.10.004

Cavanagh, P. & Anstis, S. (2013). The flash grab effect. Vision Research, 91, 8–20. doi: 10.1016/j.visres.2013.07.007

Cavanagh, P., Anstis, S., Lisi, M., Wexler, M., Maechler, M., ’t Hart, B. M., Shams-Ahmar, M., Saleki, S. (2022). Exploring the frame effect. Journal of Vision, 22(12):5. doi: 10.1167/jov.22.12.5.

Chung, Y. H., Störmer, V. S. (2023). Unveiling the time course of visual stabilization through human electrophysiology. iScience, 26(6), 106800. doi: 10.1016/j.isci.2023.106800.

Dewey, J. A., & Carr, T. H. (2013). Predictable and self-initiated visual motion is judged to be slower than computer generated motion. Consciousness and Cognition, 22(3), 987–995.

Duncker, K. (1929). Über induzierte Bewegung (Ein Beitrag zur Theorie optisch wahrgenommener Bewegung). Psychologische Forschung, 12, 180–259. doi: 10.1007/BF02409210. Abridged and translated (1938) as “Induced Motion” in *Source Book of Gestalt Psychology*, edited and translated by Ellis, W. D.London: Routledge and Kegan Paul. pp 161–172.

Johansson, G. (1950). Configurations in the perception of velocity. Acta Psychologica, 7, 25–79. doi: 10.1016/0001-6918(50)90003-5

Kolers, P.A. (1972). Aspects of Motion Perception. Oxford: Pergamon Press. doi: 10.1016/C2013-0-05617-4

Luke, S.G. (2017). Evaluating significance in linear mixed-effects models in R. Behavior Research Methods, 49, 1494–1502. doi: 10.3758/s13428-016-0809-y

MacLeod, D. I. A., Cavanagh, P., & Anstis, S. (2024). Contribution of low-level motion to position shifts. Journal of Vision, 24(8):13, 1–9, 10.1167/jov.24.8.13.

Maruya, K., Yang, E., Blake, R. (2007). Voluntary action influences visual competition. Psychological Science, 18 (12), 1090–1098. doi: 10.1111/j.1467-9280.2007.02030.x

Matin, L., & Fox, C. R. (1989). Visually perceived eye level and perceived elevation of objects: linearly additive influences from visual field pitch and from gravity. Vision Research, 29(3), 315–24. doi: 10.1016/0042-6989(89)90080-1.

Morgan, M., Grant, S., Melmoth, D., & Solomon, J. A. (2015). Tilted frames of reference have similar effects on the perception of gravitational vertical and the planning of vertical saccadic eye movements. Experimental Brain Research, 233(7), 2115–25. doi: 10.1007/s00221-015-4282-0.

Newsome, L. R. (1972). Visual angle and apparent size of objects in peripheral vision. Perception & Psychophysics, 12(3), 300–304.

Nijhawan, R. (1994). Motion extrapolation in catching. Nature, 370, 256–257. doi: 10.1038/370256b0

Özkan, M., Anstis, S., ’t Hart, B. M., Wexler, M., & Cavanagh, P. (2021). Paradoxical stabilization of relative position in moving frames. Proceedings of the National Academy of Sciences, 118(25), 1–8. doi: 10.1073/pnas.2102167118

Peirce, J. W., Gray, J. R., Simpson, S., MacAskill, M. R., Höchenberger, R., Sogo, H., Kastman, & E., Lindeløv, J. (2019). PsychoPy2: experiments in behavior made easy. Behavior Research Methods, 51, 195–203. Doi: 10.3758/s13428-018-01193-y

Porac, C. (1989). Is visual illusion decrement based on selective adaptation? Perception & Psychophysics, 46(3), 279–283.

R Core Team (2025). R: A Language and Environment for Statistical Computing. https://www.R-project.org/

Roelofs, C. O. (1935). Die Optische Lokalisation. Archiv fur Augenheilkunde, 109, 395–415.

Satterthwaite, F. E. (1941). Synthesis of variance. Psychometrika, 6, 309–316. doi: 10.1007/BF02288586

Shams, M., Kohler, P.J., & Cavanagh, P. (2024). Deconstructing the frame effect. Journal of Vision, 24(11):8. doi: 10.1167/ov.24.11.8

Sinico, M., Parovel, G., Casco, C., & Anstis, S. (2009). Perceived shrinkage of motion path. Journal of Experimental Psychology: Human Perception and Performance, 35(4), 948–957. doi: 10.1037/a0014257

Skora, L. I., Seth, A. K., Scott, & R. B., (2021). Sensorimotor predictions shape reported conscious visual experience in a breaking continuous flash suppression task. Neuroscience of Consciousness, 1 (2021). doi: 10.1093/nc/niab003

Veto, P., Uhlig, M., Troje, N. F., Einhäuser, W. (2018). Cognition modulates action-to-perception transfer in ambiguous perception. Journal of Vision, 18, 5. doi: 10.1167/18.8.5

Wallach, H., Bacon, J., Schulman, P. (1978). Adaptation in motion perception: Alteration of induced motion. Perception & Psychophysics, 24(6), 509–514. doi: 10.3758/BF03198776

Warren, H. C. (1895). Sensations of rotation. Psychological Review, 2(3), 273–276. doi: 10.1037/h0074437

Wohlschläger, A. (2000). Visual motion priming by invisible actions. Vision Research, 40 (80), 925–930. doi: 10.1016/S0042-6989(99)00239-4

Wong, E., & Mack, A. (1981). Saccadic programming and perceived location. Acta Psychologica, 48, 123–131. doi: 10.1016/0001-6918(81)90054-8

